# Rye: genetic ancestry inference at biobank scale

**DOI:** 10.1101/2022.04.15.488477

**Authors:** Andrew B. Conley, Lavanya Rishishwar, Maria Ahmad, Shivam Sharma, Emily T. Norris, I. King Jordan, Leonardo Mariño-Ramírez

## Abstract

Biobank projects around the world are generating genomic data for many thousands and even millions of individuals. Computational methods are needed to handle these massive data sets, including tools for genetic ancestry (GA) inference. Current methods for GA inference are generally accurate, but they are slow and do not scale to biobank-size genomic datasets. Here we present Rye – a new algorithm for GA inference at biobank scale. We compare the accuracy and runtime performance of Rye to the widely used RFMix and ADMIXTURE programs, and we apply it to a dataset of 488,221 genome-wide variant samples from the UK Biobank. Rye infers GA based on principal component analysis (PCA) of genomic variant samples from ancestral reference populations and query individuals. The algorithm’s accuracy is powered by Metropolis-Hastings optimization and its speed is provided by non-negative least squares (NNLS) regression. Rye produces highly accurate GA estimates for three-way admixed populations – African, European, and Native American – compared to RFMix and ADMIXTURE (*R*^2^ = 0.998 – 1.00), and shows 50x runtime improvement compared to ADMIXTURE on the UK Biobank dataset. Rye analysis of UK Biobank samples demonstrates how it can be used to infer GA at different levels of relatedness. We discuss user consideration and options for the use of Rye; the program and its documentation are distributed on the GitHub repository: https://github.com/healthdisparities/rye.

## INTRODUCTION

Genetic ancestry (GA) refers to genetic similarities indicating the geographic origins of common ancestors (1,2). The GA of modern humans reflects recurrent historical patterns of migration, followed by geographical and reproductive isolation, and subsequent admixture whereby previously isolated populations come back together (3–5). GA is a characteristic of the genome, and it can be inferred based on correlated allele frequency differences that accumulate owing to the action of evolutionary forces on ancestral populations (6). GA can be characterized objectively and with precision, as a categorical or continuous variable, at the genome-wide or local level, and for different levels of relatedness. In this way, GA is distinct from the socially ascribed and more subjective categories of race and ethnicity (7,8).

Studies of GA have been widely used to illuminate the complex evolutionary history of our species (9–16). GA inference can also be used to help understand how genetic variation within and between populations contributes to health and disease. For example, the characterization of GA is crucial for the application of genome-wide association studies and polygenic risk prediction to globally diverse populations (17–19). GA can be used to help decompose genetic and environmental contributions to health disparities since it is characterized independently of the social dimensions of race and ethnicity (7,8).

There are numerous programs available for GA inference (6), including tools that characterize genome-wide ancestry (20) and local ancestry (21), along with applications for fine-scale ancestry and admixture (22). Current methods for GA inference produce accurate and reliable results, but they don’t scale well to increasingly large genomic datasets. Biobanks projects, such as the UK Biobank (23) and the NIH All of Us project (24), are generating genome-wide variant datasets for hundreds of thousands of individuals, and similarly ambitious biobank projects are underway around the world (25). Biobanks promise to revolutionize precision medicine, but their potential will not be fully realized without the development of the computational methods needed to analyze such massive datasets. GA inference methods that scale to biobank-size genomic data, while retaining the accuracy of previous generation methods, are urgently needed.

We present the program Rye as one solution to this challenge. Rye provides for rapid and accurate genome-wide GA inference on biobank-size genomic datasets, and it can be used to infer GA at different levels of evolutionary relatedness. Rye and its source code are made freely available; it is well documented, easy to install and use, relies on standard genomic variant formats, and works well with limited computational resources.

## MATERIALS AND METHODS

### Algorithm overview

The Rye algorithm infers genome-wide ancestry fractions for individual genomic variant samples by comparing principal component (PC) vectors of global reference population individuals with PC vectors of query individuals (Figure 1). Reference populations are grouped into user-defined ancestry groups, which can be assigned at varying levels of biogeographic and genetic relatedness (e.g. continental or subcontinental groups). Principal component analysis (PCA) is run on a combined variant dataset of reference and query individual samples to yield PC vectors for all individuals. PC vectors representative of each ancestry group are computed via Markov chain Monte carlo (MCMC) optimization of reference group-mean PC vectors. Finally, the optimized ancestry-representative PC vectors are compared with PC vectors of query individuals, using non-negative least squares (NNLS), to generate ancestry estimates for all individual samples, expressed as fractions of each user-defined ancestry group.

**Figure 1.**
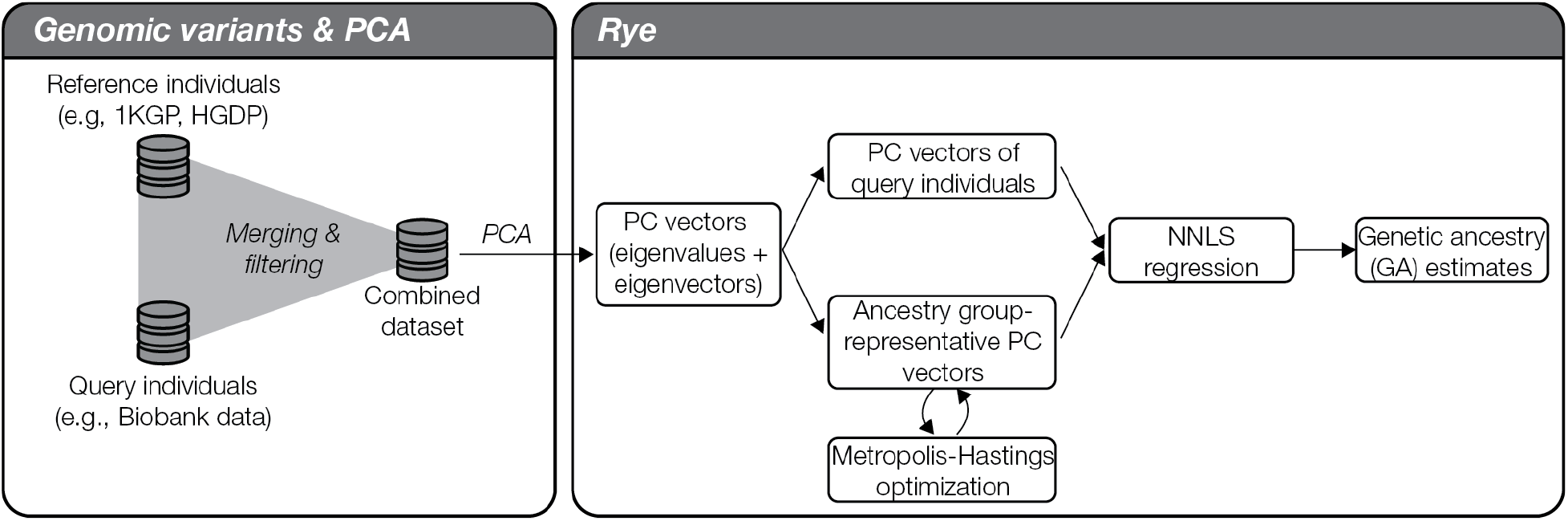
Overview of the Rye algorithm. Rye utilizes eigenvectors (PC vectors) and eigenvalues generated by PCA of reference and query individual genome-wide variant samples (left panel). Ancestry group-representative PC vectors are weighted via Metropolis-Hastings optimization of ancestry group-mean PC vectors. Non-negative least squares regression (NNLS) is used to estimate GA fractions via comparison of query individual PC vectors and the weighted ancestry group representative PC vectors.

### Algorithm workflow

A schematic illustrating the details of the Rye algorithm is shown in Supplementary Figure S1. Genetic ancestry inference with Rye is performed on a user-supplied genomic variant file that includes reference population samples and query individual samples. Ancestry inference with Rye proceeds via the following steps:

1. PCA is run on the genomic variant file to yield eigenvectors (i.e. vectors of PC values) for all reference and query individuals. PC vectors are scaled from 0 to 1 for downstream calculations.
2. Reference individuals are grouped into user-defined ancestry groups, and mean scaled PC vectors are calculated for each ancestry group.
3. Ancestry-representative PC vectors are computed from ancestry group mean PC vectors using a nested Metropolis-Hastings optimization procedure (see next section for details).
4. Optimized ancestry-representative PC vectors are used with non-negative least squares (NNLS) regression to estimate ancestry group fraction values for query individuals, with the constraint that ancestry group fraction values must sum to one. The NNLS equation below shows an example for *n* PC values and *m* reference ancestry groups, yielding *m* ancestry fractions (*β*) for a query individual PC vector.

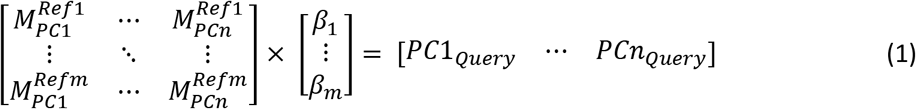

### Metropolis-Hastings optimization

A nested Metropolis-Hastings procedure is used to optimize two parameter sets for the ancestry-representative PC vectors: PC weights and shrinkage values. PC weights are used to scale the contribution of each PC to ancestry assignment, and shrinkage values are used to scale the values of each PC. PC weights are the same for all ancestry groups, whereas shrinkage values are specific to each ancestry group. PC weights are initialized using the fraction of variance explained (i.e. the eigenvalues) for each PC from the PCA, and shrinkage values are initialized uniformly. The shrinkage values are used to ensure that outlier individuals, i.e. individuals with extreme PC values compared to their ancestry group, do not bias the ancestry estimation results. The shrinkage values are used to shrink the ancestry group representative PC values towards a value of 0.5. Optimized ancestry-representative PC vectors are calculated as:

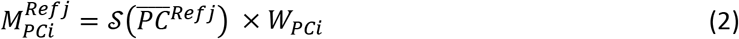

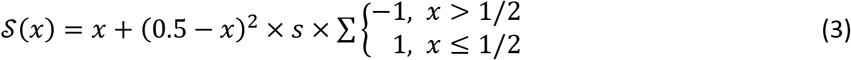

where *i* ∈ [1, *n*] PCs, *j* ∈ [1, *m*] ancestry groups, *W* is the maximum-normalized weight for *PC_i_*, and s is the shrinkage value for *PC_i_* and ancestry group *j*.

Optimization of the weight and shrinkage parameters is done using the Metropolis-Hasting algorithm, executed in a nested manner across *r* rounds, *t* attempts within each round, and *u* iterations within each attempt (Supplementary Figure S2). Rounds are executed sequentially, and attempts are launched independently within each round. Within each attempt, the Metropolis-Hastings algorithm is used to iterate over weight and shrinkage parameter values, probabilistically selecting the optimal combination of values with which to proceed after each iteration. The optimization criterion is based on NNLS prediction of group-specific ancestry values for reference individuals, as shown in equation 1 with reference individuals treated as query individuals. Reference individuals are expected to show ancestry values that maximally correspond to their group membership.

### Algorithm testing and validation

Rye was used to estimate ancestry fractions – African, European, and Native American – for three-way admixed individuals from the Americas. Rye ancestry estimates were compared against ancestry estimates obtained with the widely used RFMix (21) and ADMIXTURE programs (20). Genomic variant data were taken from the 1000 Genomes Project (1KGP) whole genome sequence data (26), and previously published set of Native American genome-wide genotypes (10). Genomic variant data from these two sources were merged and harmonized as previously described (27–29). Reference individuals were taken from African, European, and Native American populations, and query individuals were taken from Admixed American populations (Supplementary Table 1).

Algorithm accuracy was measured by comparing observed ancestry fractions for three-way admixed individuals calculated with Rye to expected ancestry fractions calculated with RFMix and ADMIXTURE using Pearson correlation (*R*^2^) and residual sum of squares (*RSS*) error. Sensitivity of the algorithm to reference sequence selection was measured using jackknife resampling with 10% of reference sequences removed in each of 10 replicates. Runtime performance was measured on a 40-core (Intel Xeon), 512GB RAM system, running on Red Hat Enterprise Linux Server release 6.10 (Santiago).

### UK Biobank

Rye was used to estimate ancestry fractions for seven regional ancestry groups – African, Central Asian, East Asian, European, Middle Eastern, Native American, and South Asian – on 488,221 participants from the UK Biobank (UKBB). UKBB participants’ genome-wide genotypes were characterized using the UKBB Axiom Array or United Kingdom BiLEVE Array as previously described (23,30). UKBB participant genome-wide genotypes were merged and harmonized with genomic variant data from global reference populations characterized as part of the 1KGP (26) and the Human Genome Diversity Project (HGDP) (31) as previously described (32,33). Reference populations were grouped into seven regional ancestry groups based on their genetic and geographic affinity. Rye ancestry estimates were compared to participants’ self-identified ethnicity (Field 21000: Ethnic background), and Rye runtime performance was compared to ADMIXTURE.

### PCA, RFmix, and ADMIXTURE

PCA was run using the FastPCA program (34) implemented in PLINK v2 (35), with data from the first 20 PCs retained for ancestry inference with Rye. RFMix was run for 12 generations in the “PopPhased”mode with a minimum node size of five and the “-use-reference-panels-in-EM” for two rounds of expectation maximization (EM). RFmix ancestry assignments were made for chromosomal regions where the RFMix ancestral certainty was at least 95%. ADMIXTURE v.1.30 was run in the unsupervised mode using default settings, with K=3 for the admixed American populations across K=3-20 for the UKBB.

## RESULTS AND DISCUSSION

### Accuracy and runtime performance

Rye was used to characterize the genetic ancestry of three-way admixed individuals from the Americas (Supplementary Table 1), and the observed Rye results were compared to expected results obtained from the widely used RFMix and ADMIXTURE programs. Rye is run using a nested optimization approach across a specified number of rounds, attempts, and iterations. Fractions of African, European, and Native American ancestry estimated for Rye and RFMix are highly correlated at both the low and high end of the numbers of rounds and iterations (Figure 2A). Similar high correlations can be seen when Rye ancestry fractions are compared to ADMXITURE estimates (Supplementary Figure S3). Higher numbers of rounds and iterations yield more accurate results, but the increase in accuracy with increasing rounds and iterations is marginal (Figure 2B). Increasing the number of rounds and iterations entails a marked decrease in runtime performance (Figure 2C). Runtime increases three-orders of magnitude at the highest numbers of rounds and iterations; nevertheless, the longest runtime is just over 20 minutes.

**Figure 2.**
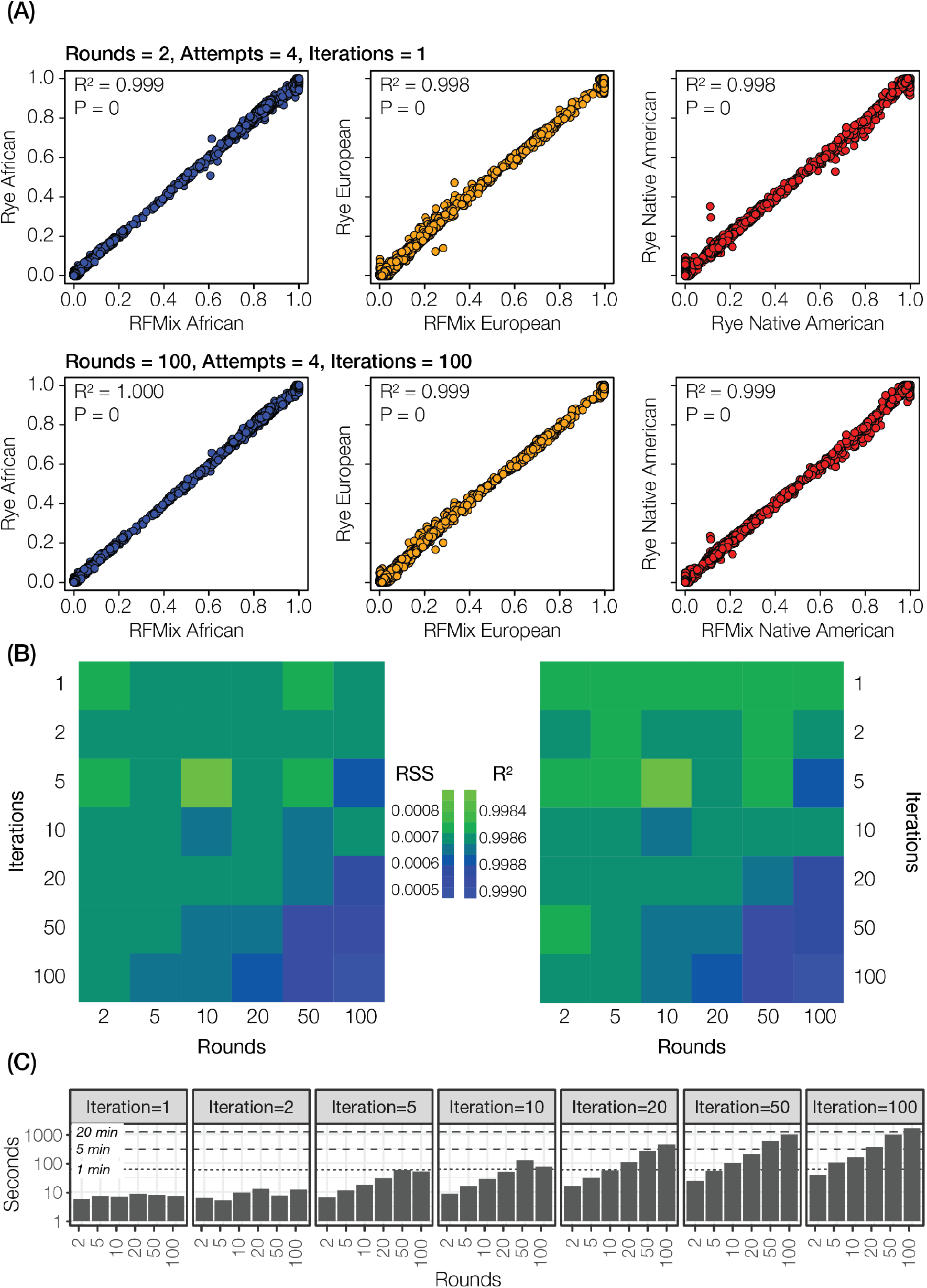
Accuracy and runtime performance. (A) GA estimates – African (blue), European (orange), and Native American (red) – are compared for Rye (y-axis) and RFMix (x-axis). (B) Accuracy of Rye measured by residual sum of squares (*RSS*) and *R*^2^ across a range of optimization rounds and iterations. (C) Runtime performance of Rye across a range of optimization rounds and iterations.

We assessed the sensitivity of Rye performance to changes in ancestry group reference samples. Jacknife resampling was used to remove 10% of reference samples across 10 replicates, and this procedure was repeated across multiple rounds and iterations (Supplementary Figure S4). Rye is relatively insensitive to changes in the composition of ancestry group reference samples. High accuracy (low RSS) is achieved even at the lowest numbers of rounds and iterations. Increasing the number rounds and iterations leads to marginal improvements in accuracy, comparable to what is seen when the full reference sample sets are used.

### Biobank scale performance

The scalability of Rye was evaluated using the UK Biobank (UKBB); genetic ancestry estimates were computed for 488,221 participants. The genetic relationship among UKBB participants and reference samples from seven regional ancestry groups, computed using PCA, are shown in Figure 3A. UKBB participants’ self-identified ethnicity are mapped onto their genetic relationships in Figure 3B. Genetic ancestry fractions for each of the seven regional ancestry groups are shown for six ethnic groups (Figure 3C). The Chinese and White ethnic groups shown the most homogenous ancestry patterns, East Asian and European respectively, whereas the Mixed and Other groups are highly diverse. The Asian ethnic group shows mostly South Asian ancestry, followed by East Asian and Middle Eastern components. The Black ethnic group shows most African ancestry followed by European and Middle Eastern components. The ancestry estimates are consistent with participants’ self-identified ethnic backgrounds, which is a second level of ethnic identity beneath the ethnic group designation (https://biobank.ctsu.ox.ac.uk/crystal/field.cgi?id=21000).

**Figure 3.**
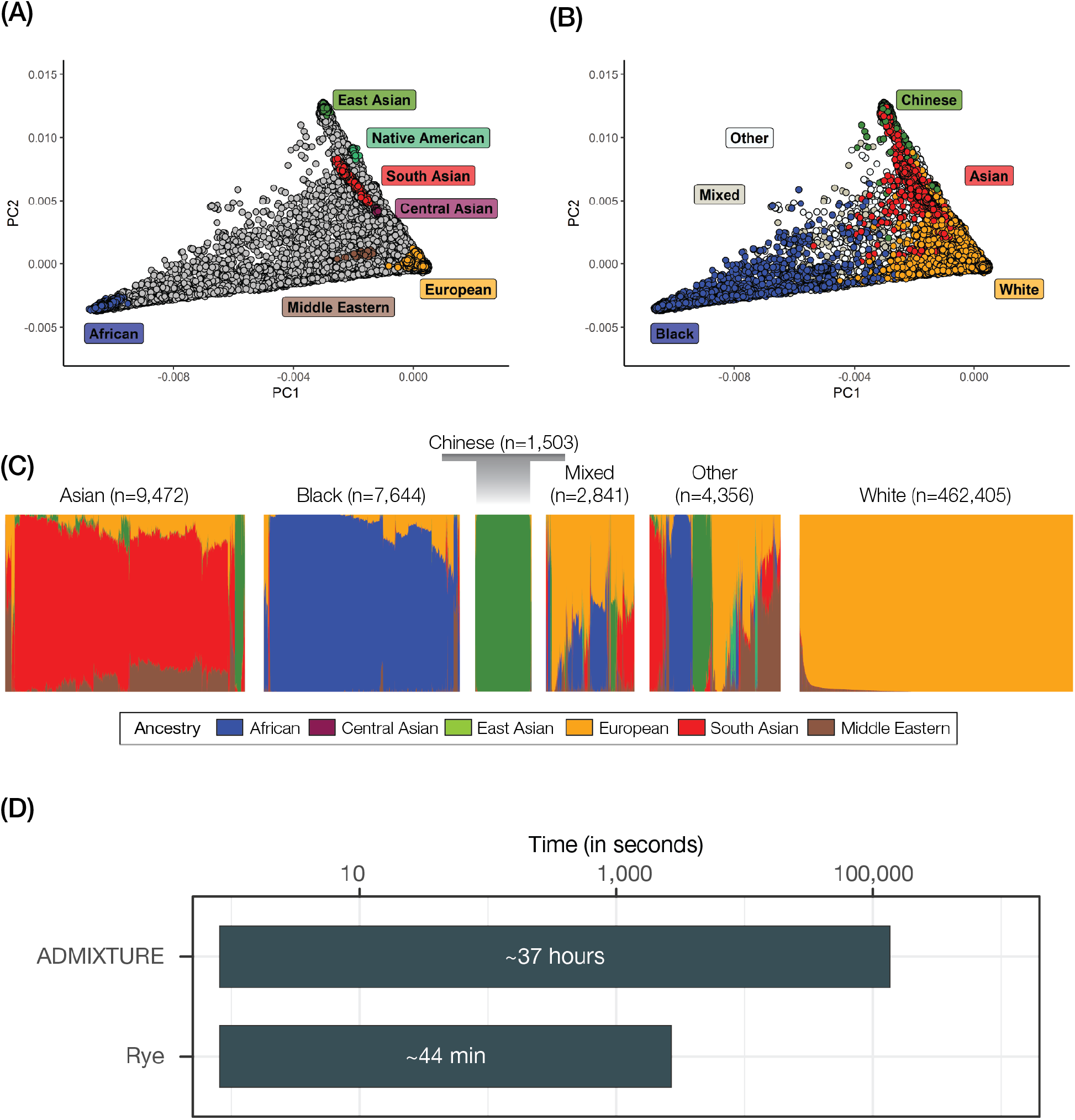
GA inference on the UK Biobank (UKBB). (A) PCA of UKBB participants (gray) and ancestry group reference samples (colored as shown). (B) PCA of UKBB participants labeled by self-identified ethnicity (colored as shown). (C) Ancestry and admixture patterns for UKBB participants, organized by self-identified ethnicity groups. Ancestry fractions (colored as shown) are indicated for each individual. The White ethnic group is not shown to scale owing to its large size; all other groups are scaled based on the number of participants. (D) Runtime comparison for ADMIXTURE and Rye, decomposed into model building (the optimization step for Rye) and GA projection steps.

Rye can also be used for GA inference at closer levels of relatedness via the delineation of more fine-scale reference ancestry groups (Figure 4). When Rye is run in this way, it reveals clearly distinct ancestry components that constitute broader East Asian, South Asian, African, and European ancestry groups. For example, UKBB participants that identify Pakistani, Indian, and Bangladeshi ethnic backgrounds, within the broader Asian ethnic group, show a gradient of distinct ancestry patterns. Similar results can be seen for African and European ancestry. African reference populations show distinct fine-scale ancestry patterns, and admixed New World African populations show largely similar African ancestry but distinct non-African admixture patterns. Rye clearly distinguishes Finnish, Russian, British, Spanish, and Italian ancestry within Europe.

**Figure 4.**
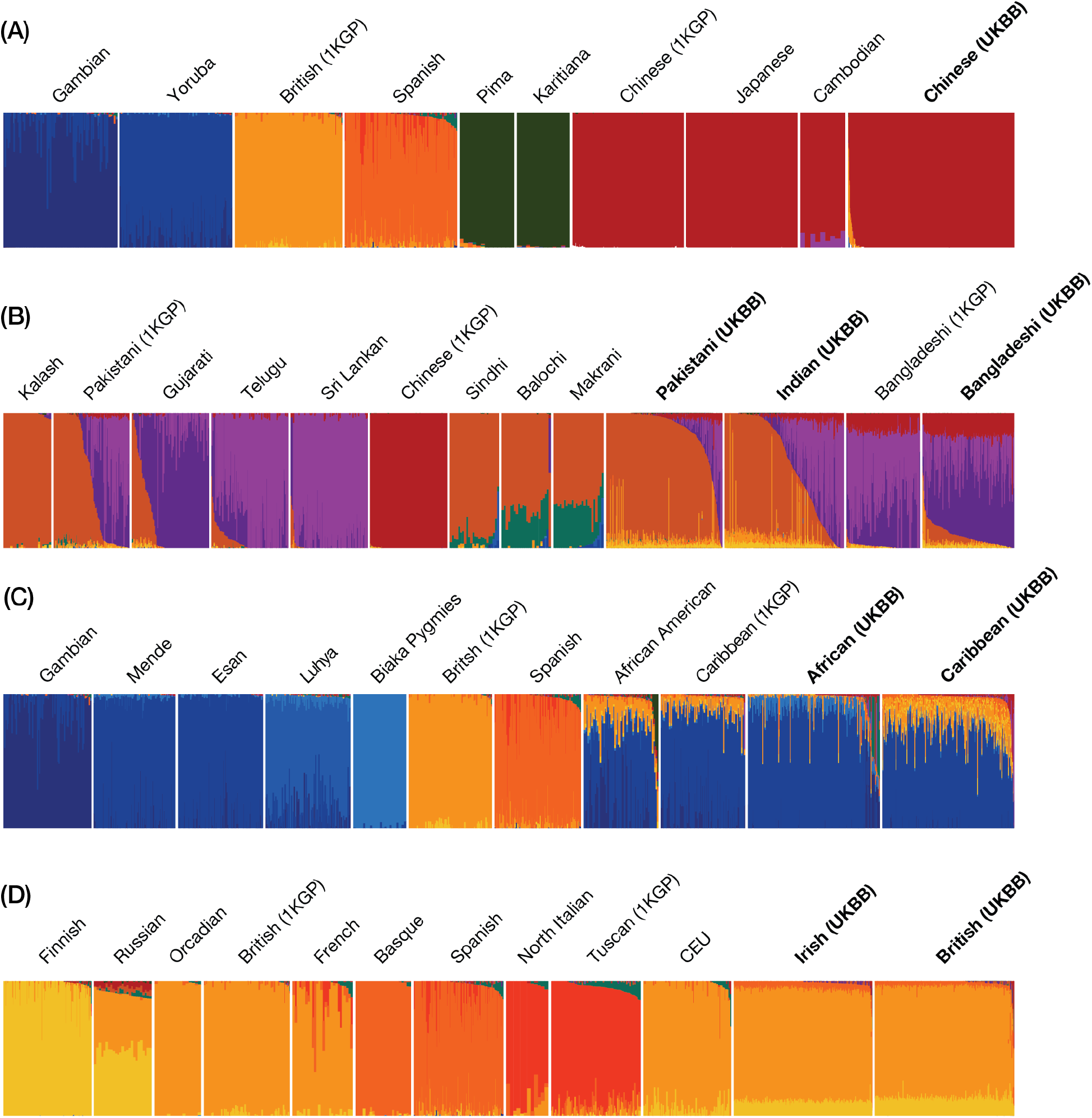
Fine-scale GA inference with Rye. Results for UKBB, 1KGP, and Native American reference and query individuals are shown. GA estimates for (A) East Asian, (B) South Asian, (C) African, and (D) European query individuals from UKBB are shown along with 1KGP and Native American reference populations.

The runtime performance of Rye on the 488,221 UKBB participants was compared to ADMIXTURE (Figure 3C). The runtime for both programs was decomposed into model building and projection phases. For Rye, model building corresponds to optimization phase and projection corresponds to the NNLS ancestry fraction calculation. Optimization was performed at the high end of Rye options, with 200 rounds and 200 iterations, to yield a conservative runtime estimate. Overall, Rye is >50x faster than ADMIXTURE: 2,687 seconds for Rye (~45 minutes) compared to 136,031 seconds for ADMIXTURE (~38 hours). Model building is ~2x faster in ADMIXTURE, but this can be attributed to the large number of rounds and iterations used for Rye optimization, which can be substantially reduced without appreciable loss of accuracy. RFMix runs prohibitively slow on a dataset of this size, and therefore its runtime performance could not be directly compared to Rye. It is estimated that RFMix would take more than two years to characterize a dataset of this size on the system used here.

### User considerations and options

Documentation and instructions for running Rye are provided on the GitHub repository: https://github.com/healthdisparities/rye. Starting with a merged reference and query genotype file, users need to run PCA and provide Rye with the output eigenvalue and eigenvector files. Samples in the eigenvector output file should include population labels in the first column and sample identifiers in the second column. Rye also requires a population to ancestry group mapping file. These three files are the only required arguments for Rye: --eigenval --eigenvec --pop2group. Other important arguments for Rye include the number of rounds, attempts, and iterations. Higher numbers for each yield more accurate ancestry estimates at the cost of slower runtime (Figure 2B). The default settings are set towards the upper end for these values, yielding the most accurate ancestry estimates. Users with large datasets or limited computational resources may considering reducing the value of these parameters. An order of magnitude time savings can be achieved in this way with little loss of accuracy (Figure 2A).

The choice of individual reference samples to be used for each ancestry group is an important consideration when using Rye. The optimization criteria for the Metropolis-Hastings algorithm assumes that reference individuals will have close to 100% ancestry for each reference group. Accordingly, the use of individuals with distinct ancestry, or admixed individuals, within the same ancestry group could impact the accuracy of ancestry estimates. Users are advised to select a subset of individuals from any given reference population, or closely related group of reference populations, that have highly similar and coherent ancestry patterns. It is not always possible to depend on population labels to choose a coherent reference individual sample set.

## Data Availability

The Rye program, its source code and documentation are freely distributed on the GitHub repository: https://github.com/healthdisparities/rye.

## Funding

ABC, LR, ETN, MA, SS and IKJ were supported by the IHRC-Georgia Tech Applied Bioinformatics Laboratory (RF383). LMR was supported by the National Institutes of Health (NIH) Distinguished Scholars Program (DSP) and the Division of Intramural Research (DIR) of the National Institute on Minority Health and Health Disparities (NIMHD) at NIH (1ZIAMD000016 and 1ZIAMD000018).

## Acknowledgements

This study was made possible by the United Kingdom Biobank application number 65206.

## Supplementary Information

### Supplementary Figures

**Supplementary Figure S1.**
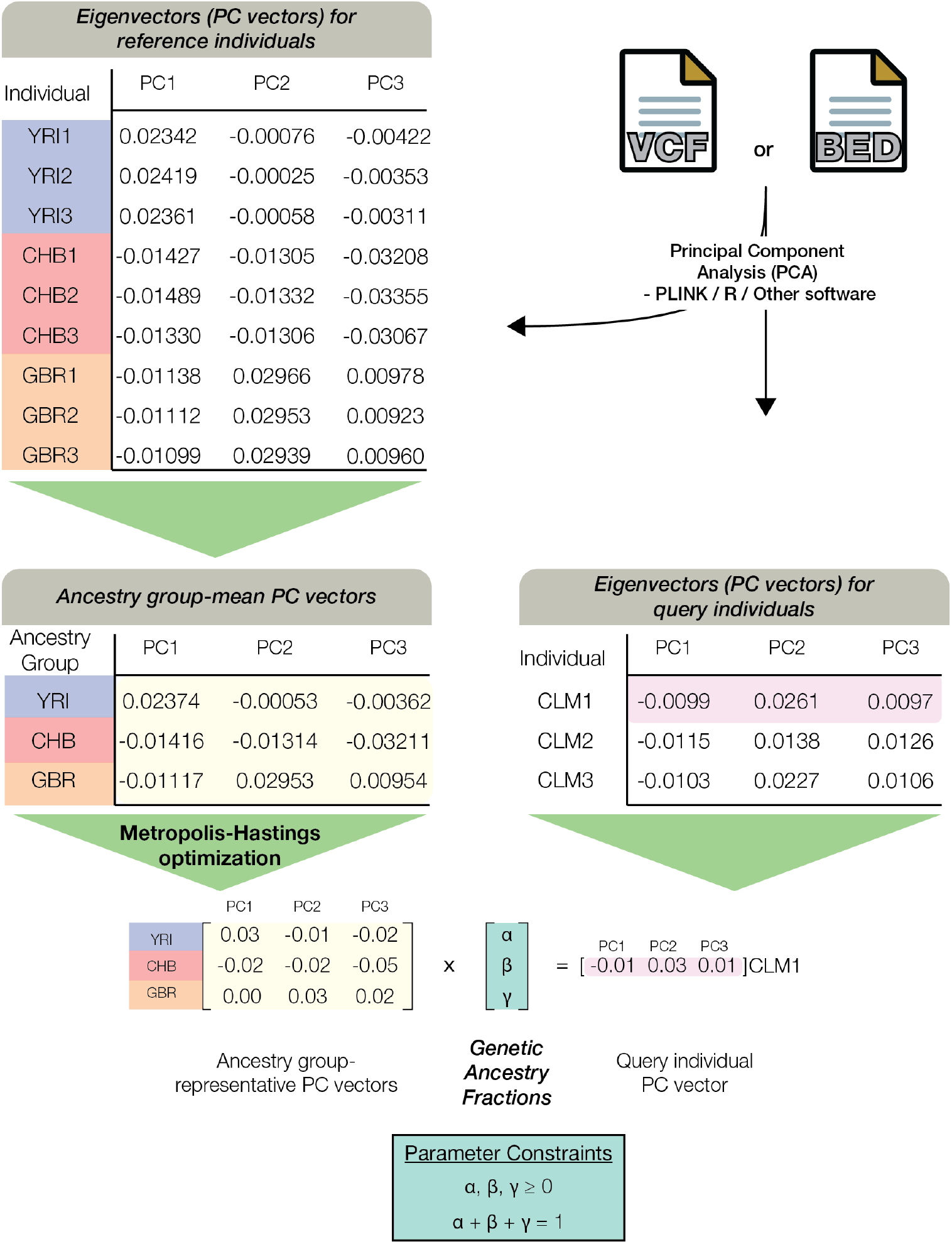
Algorithmic workflow of Rye. Rye uses PCA results to infer GA. The Rye algorithm can be divided into three broad steps. Step # 1: Input of eigenvectors (PC vectors) for each individual and mapping of reference individuals to their respective populations and ancestry groups. In this example, we consider three reference individuals each from Nigerian (YRI), Chinese (CHB), and British (GBR) ancestry groups. Step #2: Ancestry group-representative PC vectors are computed from ancestry group-mean PC vectors via Metropolis-Hastings optimization (Supplementary Figure 2). Optimization iteratively refines PC-specific weights and ancestry group-specific shrinkage values. Step 3#: Genetic ancestry (GA) fractions for each individual are calculated using non-negative least square (NNLS) regression.

**Supplementary Figure S2.**
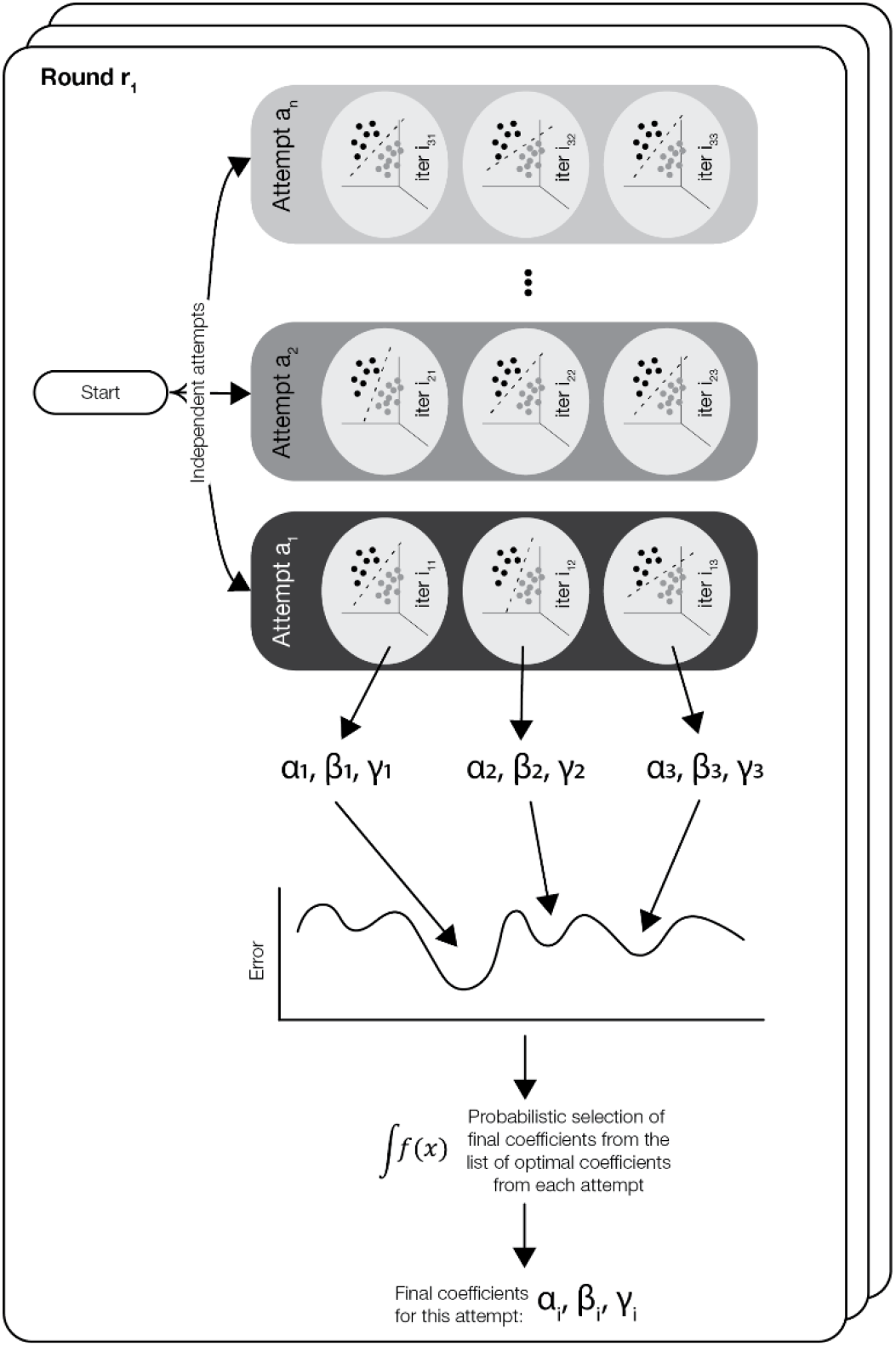
Metropolis-Hastings optimization approach implemented within Rye. Rye adopts a heuristic, three-level approach for quickly finding optimal coefficients (i.e., genomic ancestry). The three levels within Rye are: rounds, attempts, and iterations. Each round starts with an initial set of coefficients that are passed to several independent attempts. Each attempt goes through several iterations to identify the optimal coefficients for that attempt. At the conclusion of all the attempts, a probabilistic function picks final coefficients from the list of optimal coefficients obtained from each attempt. The probabilistic function is constructed so that coefficients with better fit are more likely to be picked than coefficients with worse fits. The final coefficients are considered the end-point of a round. umber of rounds, attempts, and iterations can all be defined by the user at Rye’s command line invocation.

**Supplementary Figure S3.**
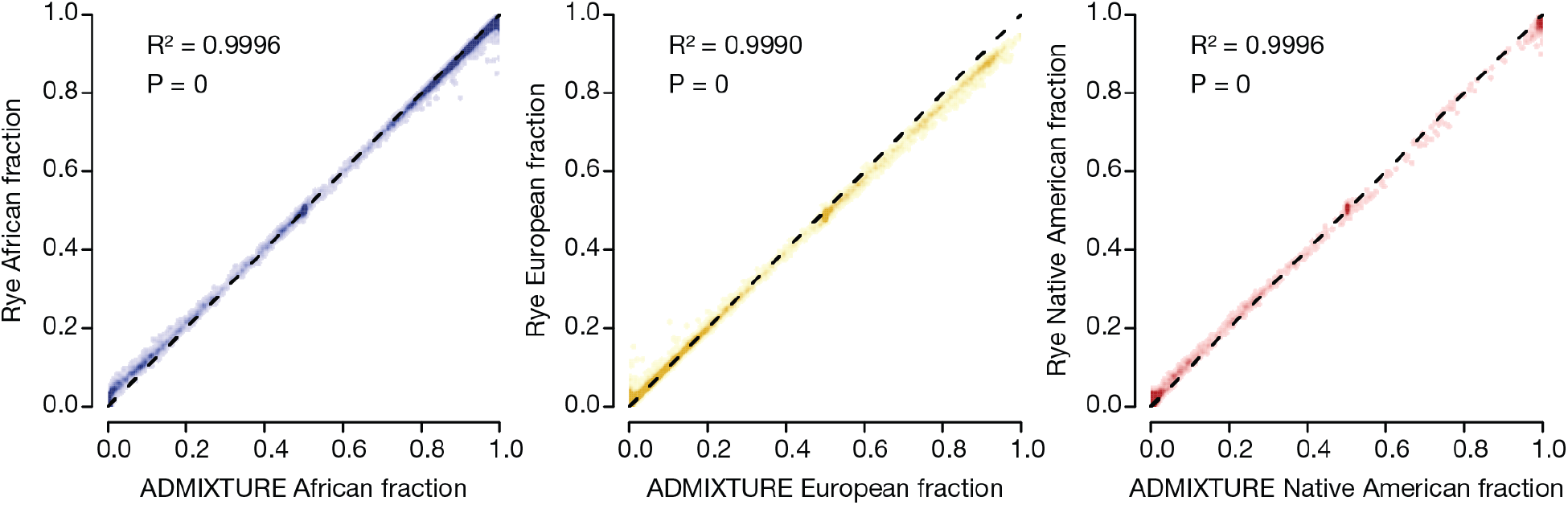
Comparison of Rye and ADMIXTURE. GA estimates – African (blue), European (orange), and Native American (red) – are regressed for Rye (y-axis) and ADMIXTURE (x-axis).

**Supplementary Figure S4.**
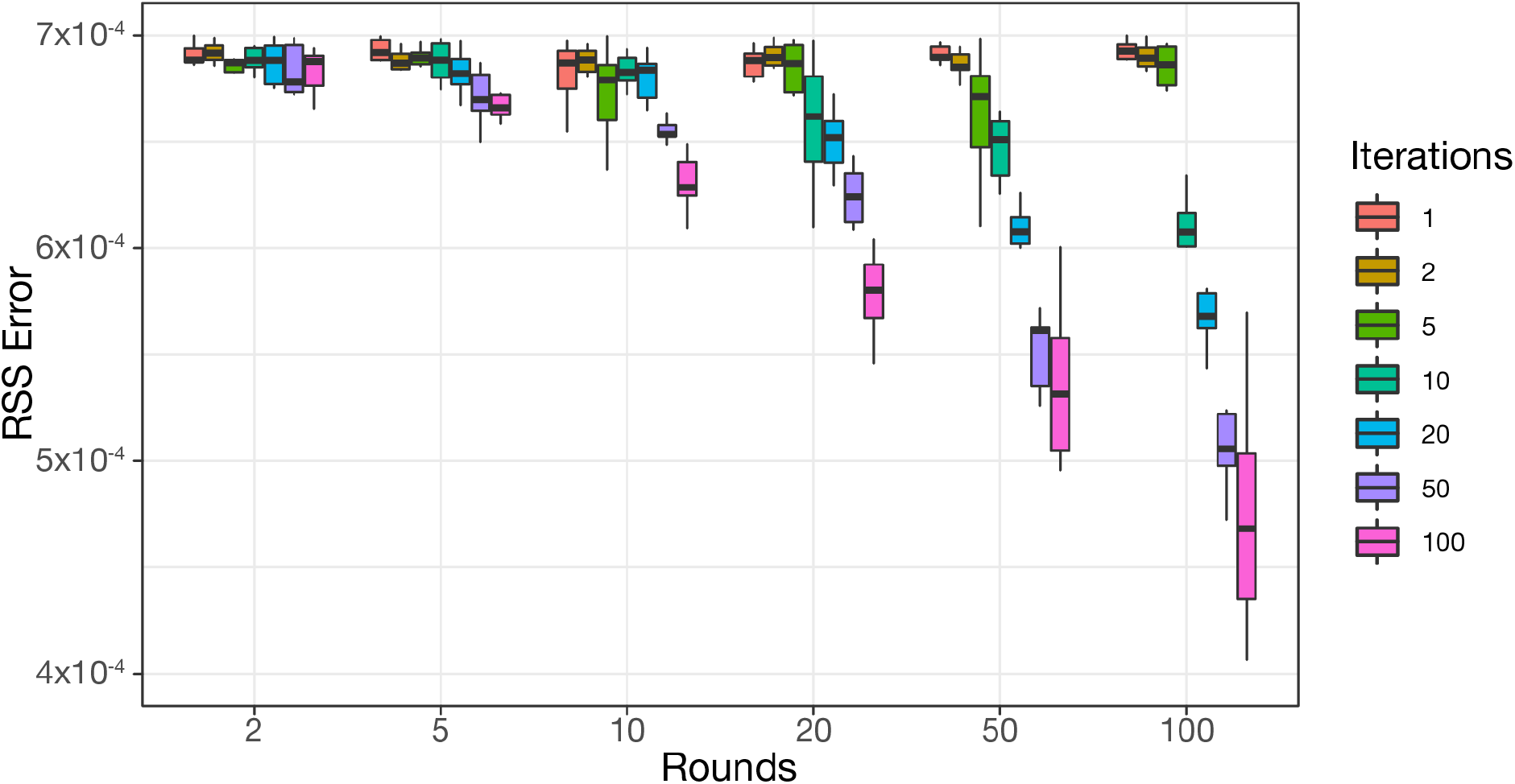
Sensitivity analysis. Rye accuracy is shown as residual sum of squares (RSS) error compared to RFMix, based on jackknife resampling analysis where 10% of reference samples were removed for 10 replicates. Results are shown across a range of rounds and iterations.

### Supplementary Tables

**Table 1.** List of populations analyzed in this study. For each population, the supergroup, number of individuals, source, and population type is indicated. Populations labelled as reference were used to construct the ancestry groups.

| Group | Population Name | N | Type | Source |
| --- | --- | --- | --- | --- |
| Admixed | African Caribbean in Barbados (ACB) | 96 | Query | 1KGP |
|  | African Ancestry in Southwest US (ASW) | 61 | Query | 1KGP |
|  | Colombian in Medellin, Colombia (CLM) | 94 | Query | 1KGP |
|  | Mexican Ancestry in Los Angeles, California (MXL) | 64 | Query | 1KGP |
|  | Peruvian in Lima, Peru (PEL) | 85 | Query | 1KGP |
|  | Puerto Rican in Puerto Rico (PUR) | 104 | Query | 1KGP |
| Africa | Esan in Nigeria (ESN) | 99 | Reference | 1KGP |
|  | Gambian in Western Division, The Gambia (GWD) | 113 | Reference | 1KGP |
|  | Luhya in Webuye, Kenya (LWK) | 99 | Reference | 1KGP |
|  | Mende in Sierra Leone (MSL) | 85 | Reference | 1KGP |
|  | Yoruba in Ibadan, Nigeria (YRI) | 108 | Reference | 1KGP |
| East Asia | Han Chinese in Beijing, China (CHB) | 103 | Reference | 1KGP |
|  | Japanese in Tokyo, Japan (JPT) | 104 | Reference | 1KGP |
| Europe | Utah residents with NW European ancestry (CEU) | 99 | Query | 1KGP |
|  | Finnish in Finland (FIN) | 99 | Reference | 1KGP |
|  | British in England and Scotland (GBR) | 91 | Reference | 1KGP |
|  | Iberian populations in Spain (IBS) | 107 | Reference | 1KGP |
|  | Toscani in Italy (TSI) | 107 | Reference | 1KGP |
| Native America | Aleutian | 8 | Query | Reich et al |
|  | Algonquin | 5 | Query | Reich et al |
|  | Arhuaco | 5 | Query | Reich et al |
|  | Aymara | 23 | Reference | Reich et al |
|  | Cabecar | 31 | Reference | Reich et al |
|  | Chilote | 8 | Query | Reich et al |
|  | Chipewyan | 15 | Query | Reich et al |
|  | Chono | 4 | Query | Reich et al |
|  | Cree | 4 | Query | Reich et al |
|  | Diaguita | 5 | Query | Reich et al |
|  | Embera | 5 | Reference | Reich et al |
|  | Guahibo | 6 | Reference | Reich et al |
|  | Guarani | 6 | Query | Reich et al |
|  | Guaymi | 5 | Query | Reich et al |
|  | Hulliche | 4 | Query | Reich et al |
|  | Inga | 9 | Query | Reich et al |
|  | Kaqchikel | 13 | Query | Reich et al |
|  | Karitiana | 13 | Reference | HGDP |
|  | Kogi | 4 | Query | Reich et al |
|  | Maya | 49 | Query | Reich et al |
|  | Mixe | 17 | Reference | Reich et al |
|  | Mixtec | 5 | Query | Reich et al |
|  | Naukan | 16 | Query | Reich et al |
|  | Ojibwa | 5 | Query | Reich et al |
|  | Palikur | 3 | Query | Reich et al |
|  | Piapoco | 7 | Reference | Reich et al |
|  | Pima | 33 | Reference | HGDP |
|  | Quechua | 40 | Query | Reich et al |
|  | Surui | 24 | Reference | HGDP |
|  | Tepehuano | 25 | Query | Reich et al |
|  | Teribe | 3 | Query | Reich et al |
|  | Ticuna | 6 | Query | Reich et al |
|  | Toba | 4 | Query | Reich et al |
|  | Waunana | 3 | Reference | Reich et al |
|  | Wayuu | 11 | Reference | Reich et al |
|  | Wichi | 5 | Query | Reich et al |
|  | Zapotec | 43 | Query | Reich et al |
| UKB | UK Biobank (UKB) | 488,221 | Query | Bycroft et al |

1KGP, 1000 Genomes Project (phase 3 data release) (PMID: 20981092, 23128226, 26432245); HGDP, Human Genome Diversity Project (PMID:11954565, 18288195, 18292342); Reich et al. (2012) refers to the study of Native American genetic ancestry (2012) (PMID:22801491).

## Notes

https://github.com/healthdisparities/rye

### Competing Interest Statement

The authors have declared no competing interest.

